# The Mosquito Electrocuting Trap As An Exposure-Free Method For Measuring Human Biting Rates By *Aedes* Mosquito Vectors

**DOI:** 10.1101/774596

**Authors:** Leonardo D. Ortega-López, Emilie Pondeville, Alain Kohl, Renato León, Mauro Pazmiño Betancourth, Floriane Almire, Sergio Torres-Valencia, Segundo Saldarriaga, Nozrat Mirzai, Heather M. Ferguson

## Abstract

**Background:** Entomological monitoring of *Aedes* vectors has largely relied on surveillance of larvae, pupae and non-host-seeking adults, which have been poorly correlated with human disease incidence. Exposure to mosquito-borne diseases can be more directly estimated using Human Landing Catches (HLC), although this method is not recommended for *Aedes*-borne arboviruses. We evaluated a new method previously tested with malaria vectors, the Mosquito Electrocuting Trap (MET) as an exposure-free alternative for measuring landing rates of *Aedes* mosquitoes on people. Aims were to 1) compare the MET to the BG-sentinel (BGS) trap gold standard approach for sampling host-seeking *Aedes* vectors; 2) characterize the diel activity of *Aedes* vectors and their association with microclimatic conditions.

**Methods:** The study was conducted over 12 days in Quinindé – Ecuador in May 2017. Mosquito sampling stations were set up in the peridomestic area of four houses. On each day of sampling, each house was allocated either a MET or a BGS trap, which were rotated amongst the four houses daily in a Latin square design. Mosquito abundance and microclimatic conditions were recorded hourly at each sampling station between 07:00-19:00 hours to assess variation between vector abundance, trapping methods, and environmental conditions. All *Aedes aegypti* females were tested for the presence of Zika (ZIKV), dengue (DENV) and chikungunya (CHIKV) viruses.

**Results:** A higher number of *Ae. aegypti* females were found in MET than in BGS collections, although no statistically significant differences in mean *Ae. aegypti* abundance between trapping methods were found. Both trapping methods indicated female *Ae. aegypti* had bimodal patterns of host seeking, being highest during early morning and late afternoon hours. Mean *Ae. aegypti* daily abundance was negatively associated with daily temperature. No infection by ZIKV, DENV or CHIKV was detected in any *Aedes* mosquitoes caught by either trapping method.

**Conclusion:** We conclude the MET performs at least as well as the BGS standard, and offers the additional advantage of direct measurement of *per capita* human biting rates. If detection of arboviruses can be confirmed in MET-collected *Aedes* in future studies, this surveillance method could provide a valuable tool for surveillance and prediction on human arboviral exposure risk.

## BACKGROUND

Mosquito--borne viruses (arboviruses) are an important cause of diseases in humans and animals. In 2017, estimates suggested that mosquitoes were responsible for approximately 137 millions human arboviral infections with dengue (DENV), chikungunya (CHIKV), and Zika virus (ZIKV) being the most important [1]. Arbovirus transmission to humans depends on multiple factors that involve spatial movement and immunity of human populations [2–7], socio-economic factors and access to basic services (especially water) [8–11], and the ecology and distribution of the mosquito vectors that transmit them [12–19]. These factors combine to determine the distribution and intensity of arboviral transmission, and generate often complex and highly heterogeneous patterns of exposure and infection [20,21]. As safe and effective vaccines for DENV, CHIKV and ZIKV viruses are not yet available [22–27], control of the *Aedes* mosquito vectors remains a primary strategy for reducing transmission [28–30].

Knowledge of where and when humans are at greatest risk of exposure to infected mosquito bites is vital for prediction of transmission intensity and effective deployment of vector control [31–34]. In the case of malaria, this information is used to estimate a time or site-specific “Entomological Inoculation Rate” (EIR); defined as the number of infected mosquito bites a person is expected to receive. This metric is usually derived from conducting Human Landing Catches (HLCs); a method in which a participant collects and counts the number of mosquito vectors landing on them over a given sampling period, then the sample is tested for the presence of a pathogen [35]. By providing a direct estimate of human exposure, the HLC provides sensitive predictions of malaria transmission [32,36–40]. However, this method raises ethical concerns due to the requirement for human participants to expose themselves to potentially infectious mosquito bites [41]. In the case of malaria, this risk can be minimized by providing participants with prophylaxis [42]. However, such remediation is not possible for arboviruses where often no prophylaxis is available, and therefore HLCs are not recommended for the surveillance *of Aedes*-borne arboviruses [43,44].

Standard entomological monitoring for *Aedes* vectors is usually based on “exposure-free” surveillance of larvae or non-biting adults. This includes surveys of larvae or pupae in water containers [45–49], and collection of adult mosquitoes resting inside and/or around houses to indirectly estimate human-vector contact rates [45,50–53]. While such surveillance methods are useful for confirming vector abundance and distribution, they are poor predictors of epidemiological outcomes such as disease incidence and outbreak potential [54,55]. Consequently there is a need for vector sampling methods that can provide more reliable entomological indicators of arboviral transmission.

Human exposure to arboviral infection is likely best assessed by surveillance of “host seeking” (human-biting) *Aedes* mosquitoes. Several methods have used to sample host seeking *Aedes* including a variety of fan-operated traps that use visual attraction cues (*e.g.* Fay [56], the Fay-Prince trap [57], the black cylinder suction trap [58], duplex cone trap [59]) and lure-based traps. For the latter, artificial odours and attractants have been developed and tested for use in traps such as kairomone blends [60–67], BG-Lure^®^ cartridges [68–70], and carbon dioxide (CO_2_) [71]. Additionally other trapping methods have been developed that use live hosts as lures (*e.g.* animal-baited traps [72] and human-baited traps [85–87).]. Only a few studies have directly compared such alternative trapping methods against the HLC with most being outperformed by the latter [74,75]. Out of all these methods, the BG-sentinel (BGS) trap has been demonstrated as one of the most effective and logistically feasible [76–81], and thus often considered a gold standard for *Aedes* surveillance [82,83]. In a range of trap evaluation studies, the BGS outperformed other methods for *Aedes* vectors with the exception of the HLC [84]. Despite these advantages of the BGS, its ability to accurately reflect the biting rates experienced by one person remains unclear. Consequently, there is still a need for a safe alternative for direct assessment of human biting rates.

Recently, a new Mosquito Electrocuting Trap (MET) was developed as an exposure-free alternative to the HLC for sampling malaria vectors [98–100]. Similar to the HLC, this sampling method also uses human participants to lure mosquito vectors and trap them. However, the MET provides participants with full protection from mosquito bites so that no exposure is required. The MET consists of four squared-shaped electrocuting surfaces that are assembled around the legs of a host, with the rest of their body being protected by netting. Host-seeking mosquitoes are attracted towards the host by odour and heat cues as normal, but are intercepted and killed before landing. In previous trials in Tanzania, the MET matched the performance of the HLC for sampling malaria vectors in rural and urban settings [85–87]. This trap has also been used to assess host preference by baiting with human and livestock hosts [87], although it has not yet been evaluated for sampling *Aedes* vectors. If successful in this context, the MET could significantly improve ability to monitor and predict arboviral transmission by facilitating an exposure-free direct estimation of EIR.

This study reports the first evaluation of METs for sampling host-seeking *Aedes* vectors in a hotspot of DENV and ZIKV transmission in coastal region of Ecuador. This region is endemic for such arboviral diseases and has accounted for most of the cases reported in Ecuador. For instance, during the CHIKV outbreak in 2015, a total of 33,625 cases were reported in Ecuador, from which 96.02% was reported in the coastal region [88]. A similar pattern occurred during the ZIKV outbreak in 2016 and 2017, where approximately 98.49% of the cases were reported in this region from a total of 5,303 cases [89,90]. DENV has been reported every year in high numbers and considering 2016 and 2017, 84.78% of cases came from the coastal region from a total of 25,537 cases [90,91].

The objectives of this study were to: (1) evaluate the performance of the MET relative to the BGS trap for sampling host-seeking *Ae. aegypti* and other mosquitoes in the study area; and (2) use the MET to characterize the biting time of *Ae. aegypti* and other relevant mosquito species and their association with microclimatic conditions.

In addition, we took the opportunity to test for the presence of arboviruses in the collected *Aedes* females by both trapping methods to investigate arboviral transmission in the local area.

## METHODS

### Location and Time of the Study

This study was conducted in the neighbourhood of “Los Higuerones” (0°19′34″N, 79°28′02″W, 78 m.a.s.l), located in the city of Quinindé (Rosa Zárate) – Ecuador. This neighbourhood is located in an urban setting dominated by small, closely packed houses (Figure 1a), bordering the eastern side with the Blanco river (Figure 1b). Quinindé is located in the province of Esmeraldas, the northernmost province in the coastal region of Ecuador. During the 2015 outbreak of CHIKV, this province accounted with the highest disease burden in the country, with a total of 10,477 cases [88]. While for DENV, during 2016, Quinindé alone accounted for 52% of the cases within Esmeraldas province, with a total of 689 cases out of a total of 1,319. In 2017, the number of DENV cases in Quinindé was much lower compared with 2016, where only 87 cases were reported out of 334 in the province of Esmeraldas. Although there is a permanent incidence of arbovirus cases along the year, a higher incidence is usually reported during the first half of the year [11].

**Figure 1.**
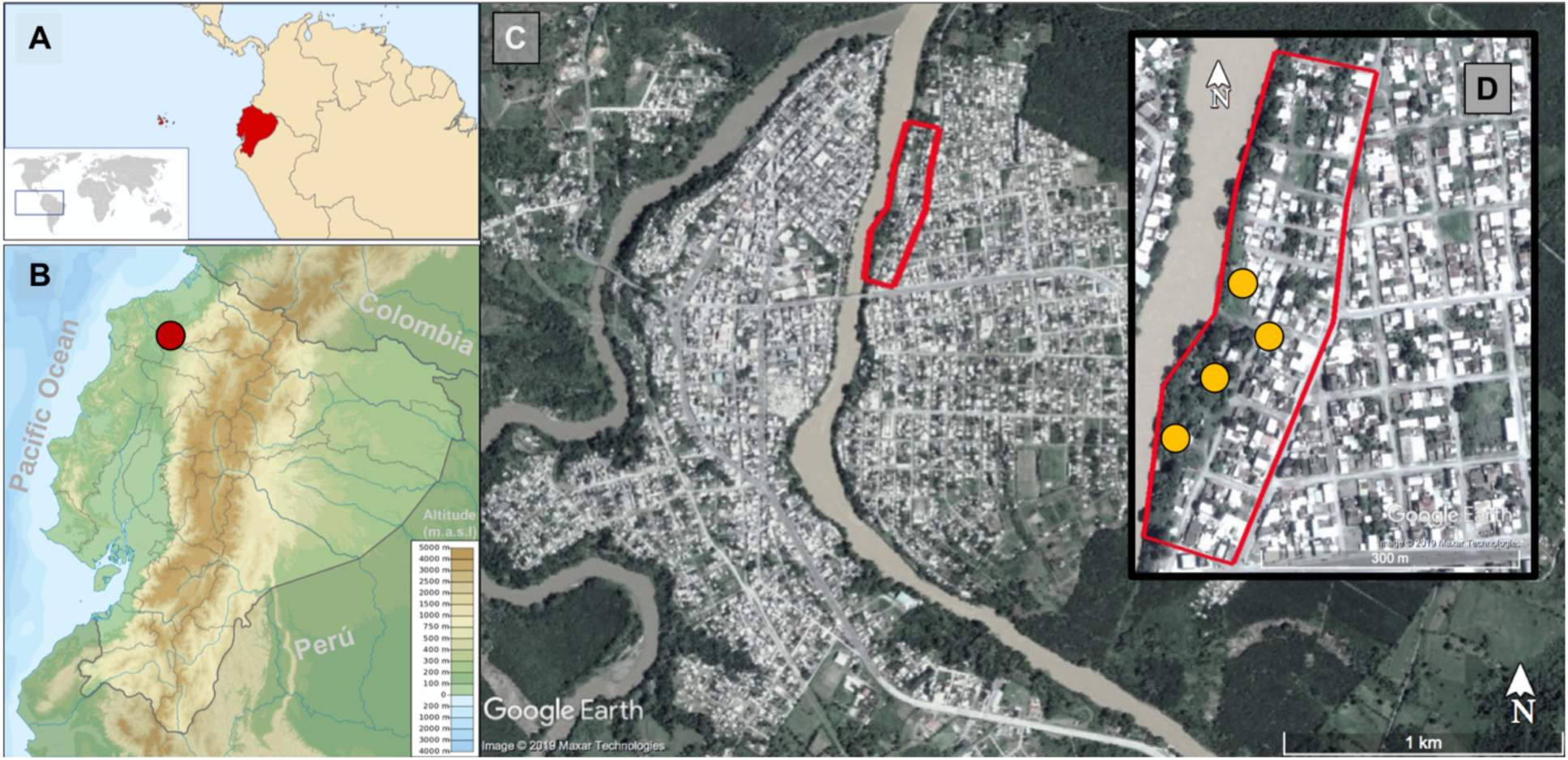
View of the urban area of the city of Quinindé. (a) Location of Ecuador in the Americas highlighted in red (Taken from [92]). (b) Location of the city of Quinindé in the Pacific Coastal region, spotted by the red circle. (c) City of Quinindé showing Los Higuerones neighbourhood enclosed by the red line. (d) Enlarged view of Los Higuerones with the houses sampled spotted by the orange circles.

The study was carried out across 12 days in May 2017 (4^th^-12^th^, and 16-18^th^). On each day of the study, mosquito sampling was conducted over 12 hours, from 07:00 – 19:00 hours. Mosquito sampling was conducted within the peri-domestic area (garden/yard) of four households (Figure 1b). These houses were selected on the basis of being physically accessible, and having residents present and willing to participate during an initial tour of the area with a local guide. Houses were separated by approximately 90 metres from one another.

### Trapping Methods

Over the study period, host-seeking mosquitoes were sampled by two different methods as follows:

#### BG-Sentinel trap (BGS)

The BG-Sentinel^®^ trap (BioGents, Regensburg, Germany) is a white, cylinder shaped trap made of plastic with a gauze cloth covering the top and a hollow black cylinder in the top centre of the trap (Figure 2a). The trap operates with a 12-volt battery that powers an internal fan that produces inwards artificial air currents. In this study, each trap was baited with two BG-Lure^®^ cartridges and a 1.4 litre cooler bottle filled with dry ice, as it is known that CO_2_ increases the catch efficiency of BGS traps [76,77,80]. Mosquitoes are attracted towards the baited traps and then sucked through the hollow black cylinder into an internal mesh bag that can be easily removed for posterior processing.

**Figure 2.**
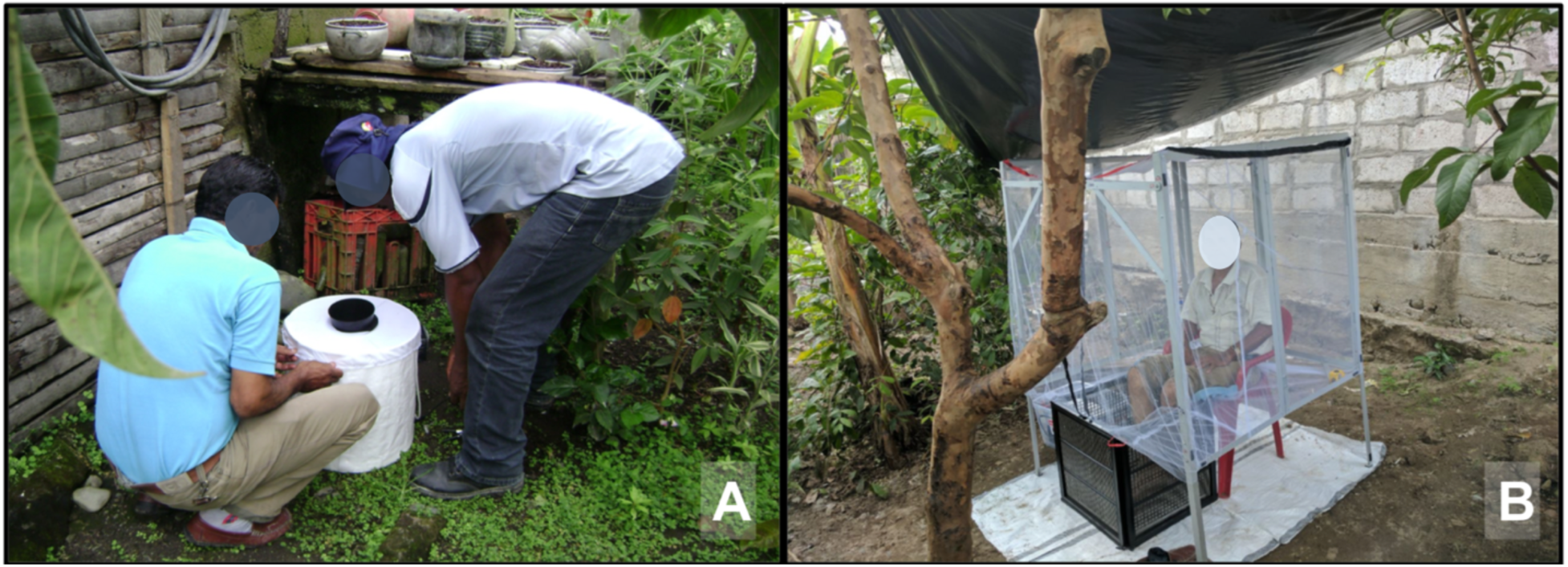
Trapping methods used in this study. (a) Typical setting up of a BGS trap. (b) Technician baiting for the MET.

#### Mosquito Electrocuting Trap (MET)

The METs used here consisted of four 30 × 30 cm panels which were assembled into a box around the lower legs of a seated person (Figure 2b). Each panel was made up of stainless steel electrified wires set within a PVC frame. The wires were positioned 5mm apart, which is close enough so that mosquitoes could not pass through without making contact. Wires were vertically arranged in parallel, alternating positive with negative. When mosquitoes try to go through, contact is made and the voltage between wires kills them.

Mosquitoes attracted towards the volunteer were intercepted and killed on contact with these panels. The MET is powered by two 12-volt batteries connected in series to a power source giving a power output of approximately 6 watts (10mA, 600 volts). An eggcrate ceiling grid panel made of plastic was placed at the inside side of each frame to provide protection to volunteers from accidentally touching the electrified wires.

As an additional accessory to the MET, a retractable aluminium frame was built to cover the rest of the volunteer’s body with untreated mosquito-proof netting. Thus volunteers were completely protected from mosquito bites during their participation in trapping. A plastic tarpaulin was erected over the MET station at a height of 2m top to protect users from direct rain and sunlight. Each MET was also set up on top of a white plastic sheet to isolate it from the ground and make it easier to see and collect shocked mosquitoes that fell onto the ground after touching the MET.

### Experimental Design

Every day of the study, four traps (two METs and two BGS traps) were set up in the peridomestic area of the four households (one trap per household) at the ground level under shade conditions. Traps were rotated among households each day, so that a different trapping method was used every consecutive day in each house. At the end of the study, this resulted in 6 days of trapping being conducted with each of the 2 methods at all houses.

MET collections were carried out by members of the research team, who were all adult men (30-50 years old). During each hour of the collection period, one member sat within the MET for 45 minutes, with the trap being turned off for the remaining 15 minutes to allow volunteers to take a break. Members of the study team took turns sitting in the trap so that different collectors lured every hour. During the 15-minute period when traps were turned off, mosquitoes were recovered from trap surfaces and the ground below using a pair of forceps, counted and placed in empty 15 ml falcon tubes; which were labelled with a unique code linked to the date, household ID, trap ID, hour period and collector ID. Tubes were stored in a cooler box of 45 L capacity filled with dry ice to kill, preserve and transport the specimens.

Each BGS was baited with two BG-Lure^®^ cartridges on each day of sampling; with lures exchanged between the two BGS traps each day to minimize bias due to differential lure efficiency. BGS traps were further baited with carbon dioxide by adding one 1.2 L Coleman^®^ polyethylene cooler bottle filled with dry ice. Dry ice containers were topped up every day. Like the MET, BGS sampling was conducted for 45 minutes of each sampling hour, with mosquito collection bags being checked and emptied during 15 minute break periods. Mosquitoes from BGS collection bags were emptied into pre-labelled plastic bags and transferred into a cooler box with dry ice to kill and preserve the mosquitoes.

Temperature and relative humidity data were collected every 10 minutes at each mosquito sampling point using TinyTag^®^ Plus 2 TGP-4500 (Gemini Co., UK) data loggers. Data loggers at the BGS sampling stations were tied and hung inside each of the traps, and loggers at MET sampling points were placed on top of the bottom border of the netting frame, next to the MET.

### Morphological Analysis

Mosquitoes collected in the field were transported to the Medical Entomology and Tropical Medicine Laboratory of the San Francisco de Quito University (LEMMT-USFQ) in cooler boxes filled with dry ice. At LEMMT-USFQ, mosquitoes were morphologically identified using taxonomical keys [93–95], counted and sorted into different cryo-vials according to date, household, trap type, hour of collection, species, sex and physiological status of females (blood fed/gravid and non-blood fed). All female *Ae. aegypti* specimens were retained for subsequent molecular analysis to test for the presence of ZIKV, DENV and CHIKV. These *Ae. aegypti* samples were grouped into pools of a maximum of 5 individuals.

### Molecular Detection of Arboviruses

All pools of female *Ae. aegypti* specimens were screened for the presence of CHIKV, DENV and ZIKV. Details on the RNA extraction, reverse-transcription and PCR procedures are given in the Supplementary File 1.

### Data Analysis

Statistical analyses were performed in R 3.5.0 and R Studio 1.1.419. Generalized Linear Mixed Models (GLMM) were used to investigate variation in the abundance of host-seeking mosquitoes (per day and per hour) using package lme4 [96]. As mosquito abundance data was overdispersed, all models were fitted with a negative binomial distribution. For all response variables of interest as described below, model selection was carried out through a process of backward stepwise elimination from a maximal model using Likelihood Ratio Tests (LRT) [97].

Statistical analysis was performed for *Ae. aegypti*, and *Culex quinquefasciatus* as the latter was the only other mosquito species found in high abundance in the study area. *Cx. quinquefasciatus* is a nuisance biting mosquito and also a known vector of West Nile Virus (WNV) [98].

The BGS traps functioned continuously across all days and sampling hours. However, the METs stopped running during some sampling hours; generally under conditions of very high humidity due to rainfall which resulted in dampness on the traps and some temporary short circuiting. When these malfunctions occurred, the damaged traps were turned off and repaired. This resulted in variation in the total number of hours sampled with each trapping method (MET: 229 hours, BGS: 270 hours). This variation in sampling effort was accounted for in the statistical analysis. Days having less than 9 hours were excluded from the analysis.

Four models were built to assess variation in the abundance of each mosquito species and sex combination respectively. For each of these four response variables, a maximal model was constructed that included the fixed explanatory variables of sampling effort (total number of hours of collection), trap type (MET or BGS), daily mean relative humidity (%RH), and daily mean temperature (°C). In addition, the interaction between daily mean temperature with relative humidity was also included. Sampling day (1 through 12), household ID, trap ID and attractant ID (BG-Lure cartridge ID or MET volunteers ID) were included as random effects.

Mosquito biting activity was assessed through analysis of variation in the mean number of females (*Ae. aegypti* and *Cx. quinquefasciatus*) caught per hour. Here, each mosquito species was analysed separately. Each model included explanatory variables of trap type (MET or BGS), sampling hour, mean temperature (°C) per hour, mean relative humidity (%RH) per hour, and the interaction between hourly temperature and relative humidity. Sampling hour was defined as a continuous variable recoding the first hour of trapping (7-8 am) into 1, and increasing “hour” by one digit for each subsequent hour until 12 (17-18 hrs). Sampling hour was fit both as a linear and quadratic term; with the latter being used to test for peaks in biting time as have been previously reported for these mosquito species [99]. In addition, sampling day, trap ID, cluster ID, household ID (nested within cluster ID) and attractant ID (BG-Lure cartridge ID or MET volunteer ID) were fitted as random effects.

## RESULTS

### Mosquito Species and Abundance

During the 12 day-experiment, a total of five mosquito species were collected by both trapping methods (Table 1). *Cx. quinquefasciatus* was the most abundant species (78.6%) followed by *Ae. aegypti* (15.63%), and small number of *Aedes angustivittatus* (2.69%), *Limatus durhami* (2.33%,) and *Psorophora ferox* (0.15%). A small proportion of mosquitoes could not be identified (0.51%, Table 1). Overall, more mosquitoes were collected with the BGS trap (60.77%) than with the MET (39.23%), but the numbers of *Ae. aegypti* were relatively similar (Table 1).

**Table 1.** Abundance of mosquito species collected by MET and BGS traps. Mosquito species abundances are split by sex and feeding status of females. The total sampling effort with the two METs was 229 hours, while for BGS traps was 270 hours over the 12 days of sampling.

|  | **Mosquito Electrocuting Trap**<br>**(MET)** |  |  |  | **BG-Sentinel (BGS) trap** |  |  |  |  |
| --- | --- | --- | --- | --- | --- | --- | --- | --- | --- |
| **Species** | ♂ | ♀<br>unfed | ♀<br>fed | **Total** | ♂ | ♀<br>unfed | ♀<br>fed | **Total** | **Grand**<br>**total** |
| *Aedes aegypti* | 100 | 99 | 19 | **218** | 93 | 91 | 27 | **211** | **429** |
| *Culex quinquefasciatus* | 496 | 238 | 44 | **778** | 960 | 345 | 77 | **1382** | **2160** |
| *Aedes angustivittatus* | 4 | 38 | 6 | **48** | 0 | 24 | 2 | **26** | **74** |
| *Limatus durhami* | 0 | 22 | 0 | **22** | 0 | 42 | 0 | **42** | **64** |
| *Psorophora ferox* | 0 | 1 | 2 | **3** | 0 | 1 | 0 | **1** | **4** |
| *Unknown* | 0 | 5 | 3 | **8** | 0 | 5 | 1 | **6** | **14** |
|  | **Total MET:** |  |  | **1077** | **Total BGS trap:** |  |  | **1668** | **2745** |

In the BGS traps, some non-target insects including house flies, butterflies, crane flies, and many fruit flies were caught. No insect taxa other than mosquitoes shown in Table 1 were caught in MET collections.

The mean daily abundance of *Ae. aegypti* was approximately 2 females and 3 males for the BGS trap, and 4 females and 4 males for the MET, but no significant differences between trapping methods were found (Table 2, Figure 3a,b). The only significant predictor of daily abundance of females *Ae. aegypti* was temperature, which exhibited a negative association (Table 2, Figure 4a). Similarly, the mean daily abundance of *Cx. quinquefasciatus* females did not significantly differ between trapping methods (Table 2, Figure 3c,d), however confidence intervals (especially for males) around estimates were very large, indicating that larger sample sizes may be required to robustly test if there were differences between trap types. The number of female *Cx. quinquefasciatus* per day varied between 16 and 207; with variation being even more pronounced for males where a high of 576 was caught on one day. The daily abundance of female *Cx. quinquefasciatus* was negatively associated with daily temperature (Table 2, Figure 4b) and positively associated with the number of hours sampled in a day, while no significant differences were found in *Cx. Quinquefasciatus* regarding any covariate (Table 2).

**Figure 3.**
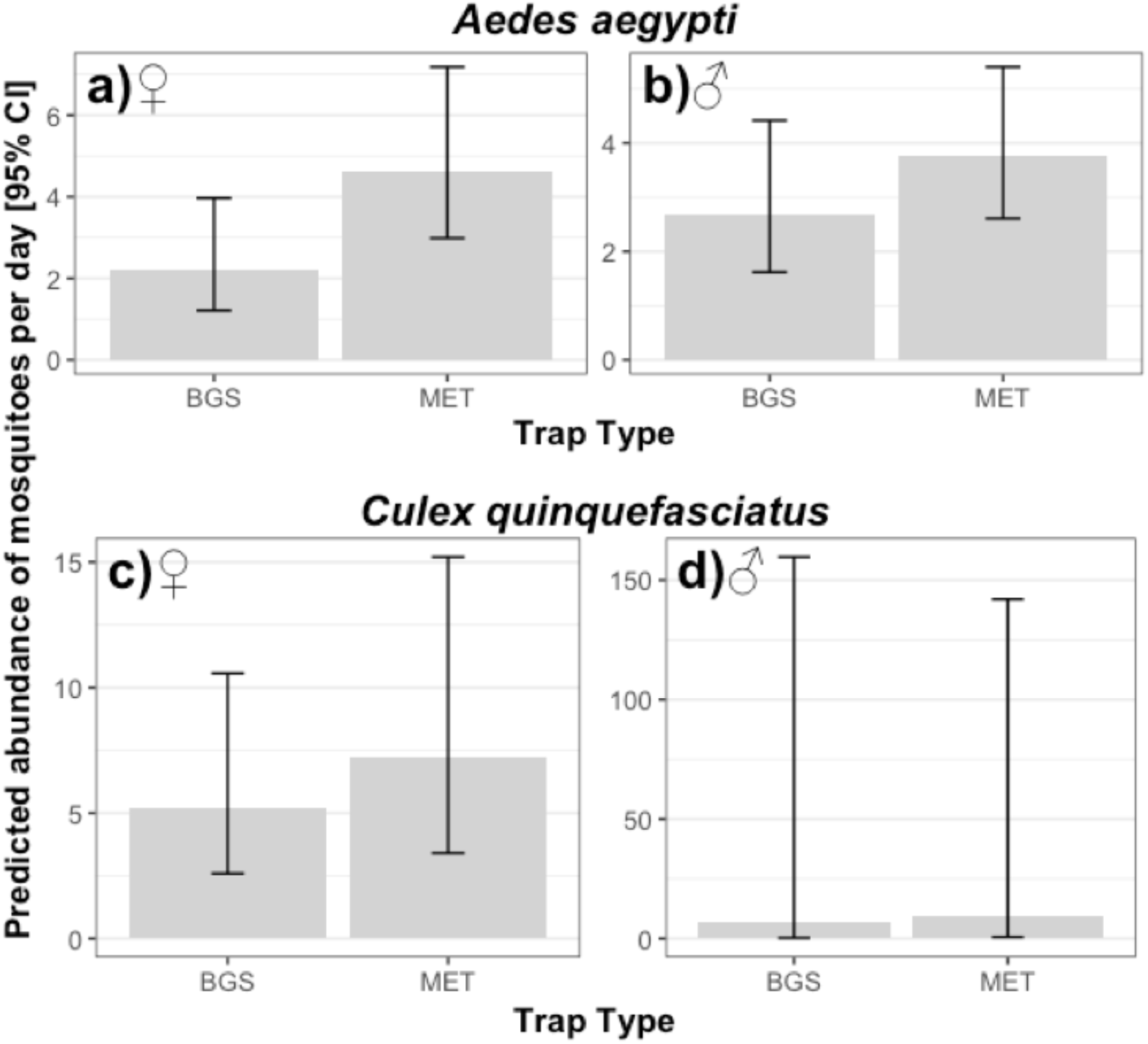
Predicted mean daily abundance of mosquitoes caught with different trapping methods. The upper panels show values for *Ae. aegypti* and the lower panels *Cx. quinquefasciatus*. Panels on the left show data for females (**♀**) and on the right for males (**♂**). Error bars indicate the Confidence Intervals (C.I.) at 95%.

**Figure 4.**
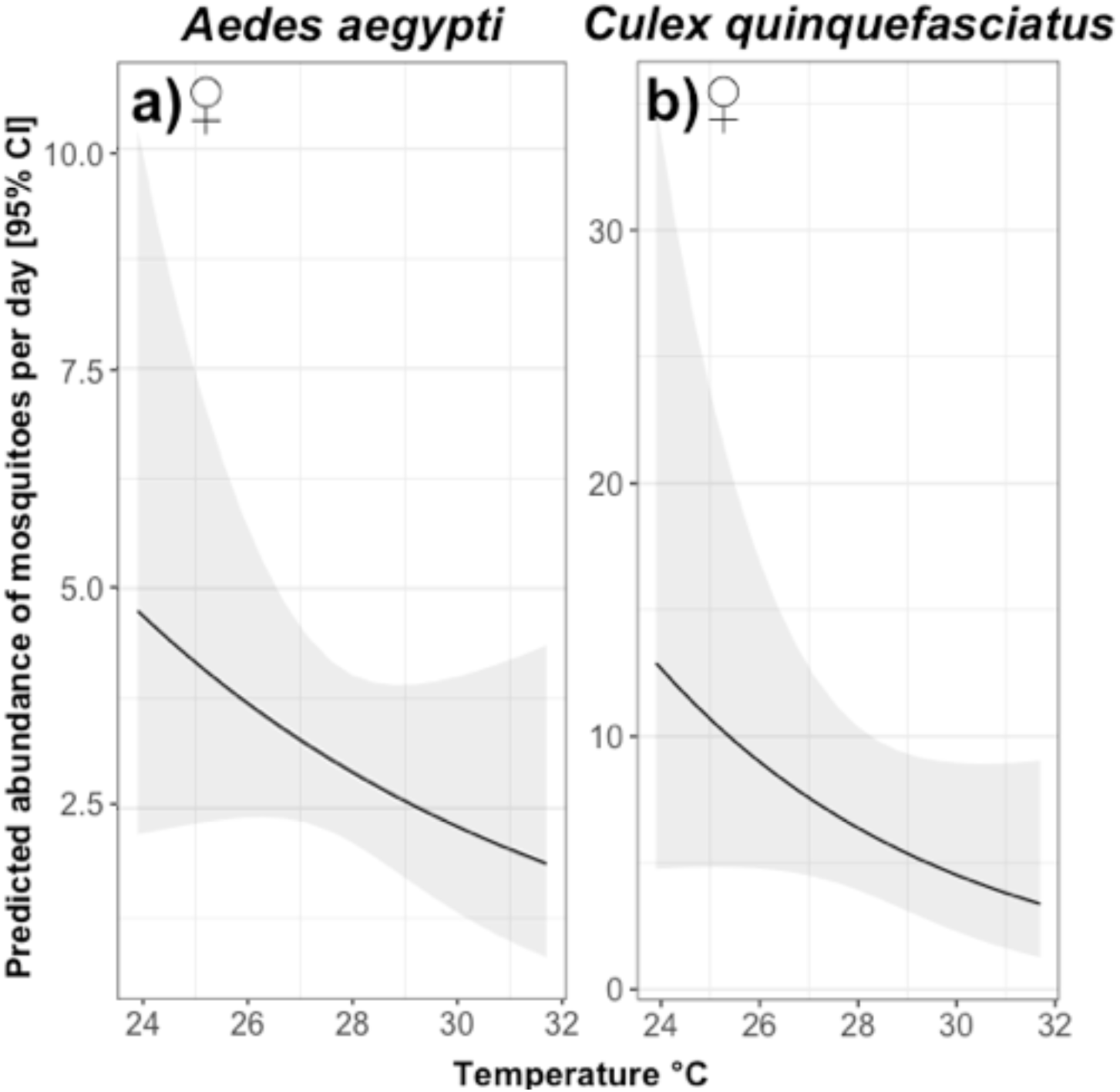
Predicted relationship between mean temperature and number of female mosquitoes collected. Panel **a)** shows *Ae. aegypti* and **b)** shows *Cx. quinquefasciatus* females. The black line indicates the mean predicted abundance and the shaded area the Confidence Intervals (C.I.) at 95%.

**Table 2.** Summary table of statistical significance of terms tested from mosquito daily abundance. Chi-square (*X^2^*), degrees of freedom (df) and *p*-values (*p*) are provided for each sex within species. Bold values with an asterisk (**\***) indicate significant terms. Fixed effects with a double S symbol (**§**) indicate the interaction term.

| Explanatory variables | *Aedes aegypti* |  |  |  |  |  | *Culex quinquefasciatus* |  |  |  |  |  |
| --- | --- | --- | --- | --- | --- | --- | --- | --- | --- | --- | --- | --- |
|  | Males ♂ |  |  | Females ♀ |  |  | Males ♂ |  |  | Females ♀ |  |  |
| | $\chi^2$ | df | $p$ | $\chi^2$ | df | $p$ | $\chi^2$ | df | $p$ | $\chi^2$ | df | $p$ |
| Sampling effort | 3.38 | 1 | 0.07 | 1.95 | 1 | 0.16 | 0.31 | 1 | 0.58 | 15.91 | 1 | **<0.001\*** |
| Trap type | 2.18 | 1 | 0.14 | 0.60 | 1 | 0.44 | 0.95 | 1 | 0.33 | 1.5 | 1 | 0.22 |
| Temperature | 0.22 | 1 | 0.64 | 4.62 | 1 | **0.03\*** | 0.06 | 1 | 0.8 | 6.86 | 1 | **<0.01\*** |
| Relative Humidity | 1.14 | 1 | 0.29 | 2.17 | 1 | 0.14 | 1.23 | 1 | 0.27 | 1.1 | 1 | 0.29 |
| Temperature :: Humidity § | 2.22 | 1 | 0.14 | 1.24 | 1 | 0.26 | 1.07 | 1 | 0.3 | 1.27 | 1 | 0.26 |

### Mosquito Biting Activity

Hourly mosquito catches recorded for BGS and METs were used to characterize the biting activity of female *Ae. aegypti* and *Cx. quinquefasciatus*. Variation in the hourly biting activity of female *Ae. aegypti* was best explained by a quadratic association between hourly mosquito abundance and time (Table 3), with activity being highest in the early mornings and late afternoon, and little activity during the middle of the day (Figure 5a). After taking this hourly variation in biting rates into account, there was no additional impact of trapping method of the number of female *Ae. aegypti* collected per hour (Table 3, Figure 6). Variation in the hourly biting activity of *Ae. aegypti* was also significantly associated with an interaction between temperature and relative humidity (Table 3). This interaction arose because the number of *Ae. aegypti* caught per hour was negatively associated with temperature under conditions of low relative humidity; but the strength of this association was lower as humidity increased (Table 3, Figure 7), although temperature and humidity were strongly associated (Figure S1).

**Table 3.** Summary table of statistical significance of terms tested for association with female mosquito hourly abundance. Chi-square (X^2^), degrees of freedom (df) and p-values are provided for females of each species. Bold values with an asterisk (**\***) indicate significant terms. Fixed effects with a double S symbol (**§**) indicate the interaction term. “N/A” indicates “not applicable” values for which single term significance was not possible because of their involvement in significant higher order terms.

| Explanatory variables | *Aedes aegypti* - Females ♀ |  |  | *Culex quinquefasciatus* - Females ♀ |  |  |
| --- | --- | --- | --- | --- | --- | --- |
| | $\chi^2$ | df | *p* | $\chi^2$ | df | *p* |
| Trap type | 0.60 | 1 | 0.44 | 7e-04 | 1 | 0.98 |
| Time (linear) | N/A | N/A | N/A | N/A | N/A | N/A |
| Time (quadratic) | 8.70 | 1 | **<0.01\*** | 142.1 | 1 | **<0.001\*** |
| Temperature | N/A | N/A | N/A | 2.07 | 1 | 0.15 |
| Relative Humidity | N/A | N/A | N/A | 0.09 | 1 | 0.77 |
| Temperature :: Humidity § | 6.60 | 1 | **0.01\*** | 0.09 | 1 | 0.76 |

**Figure 5.**
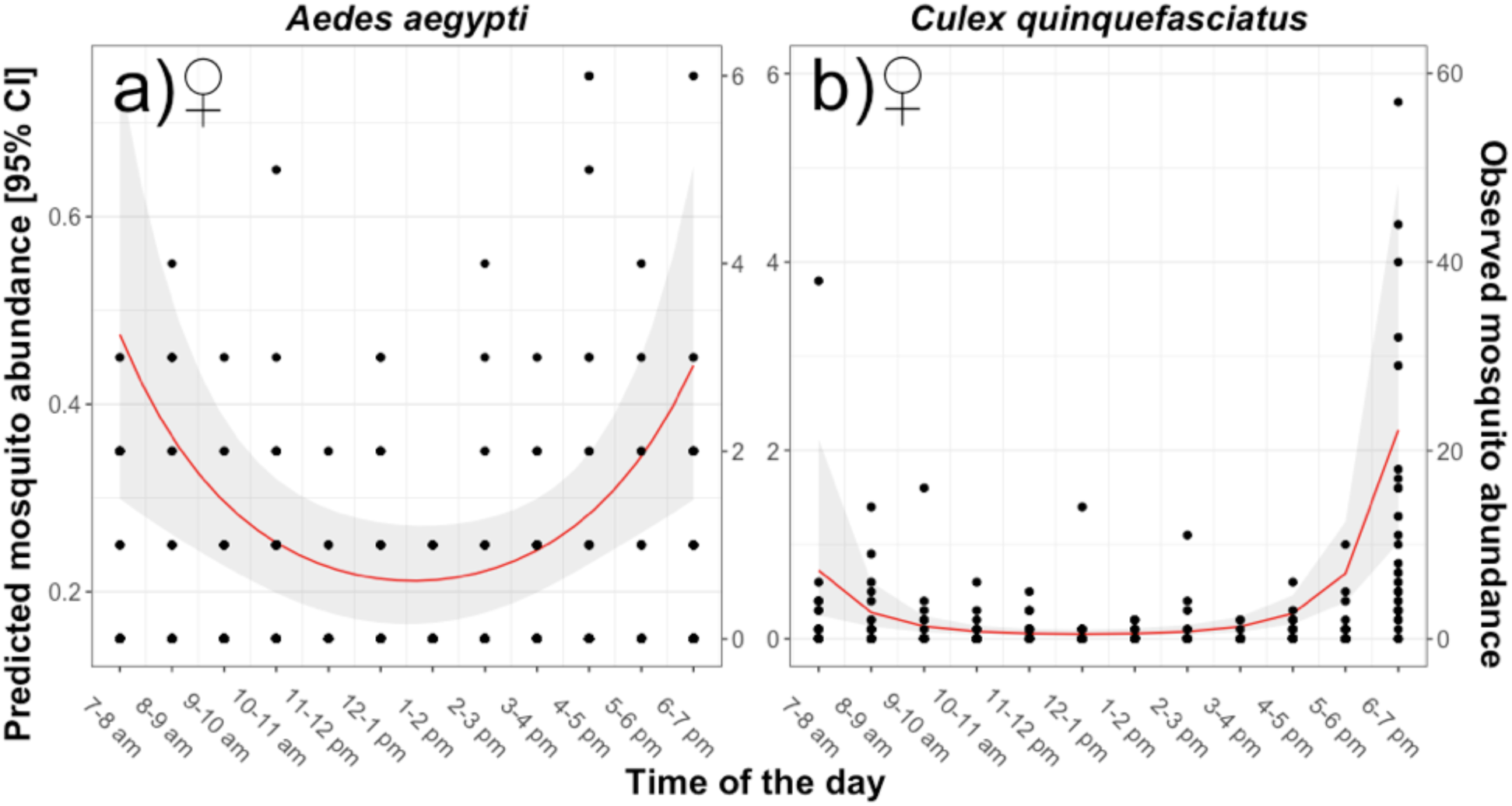
Predicted abundance of biting mosquitoes between 7:00-19:00 hrs. Panel a) indicates *Ae. aegypti* females and b) *Cx. quinquefasciatus* females. Dots represent the observed values which correspond to the right Y axis. The red line corresponds to the predicted mosquito abundance and the shaded area to the Confidence Intervals (C.I.) at 95%; both correspond to the left Y axis.

**Figure 6.**
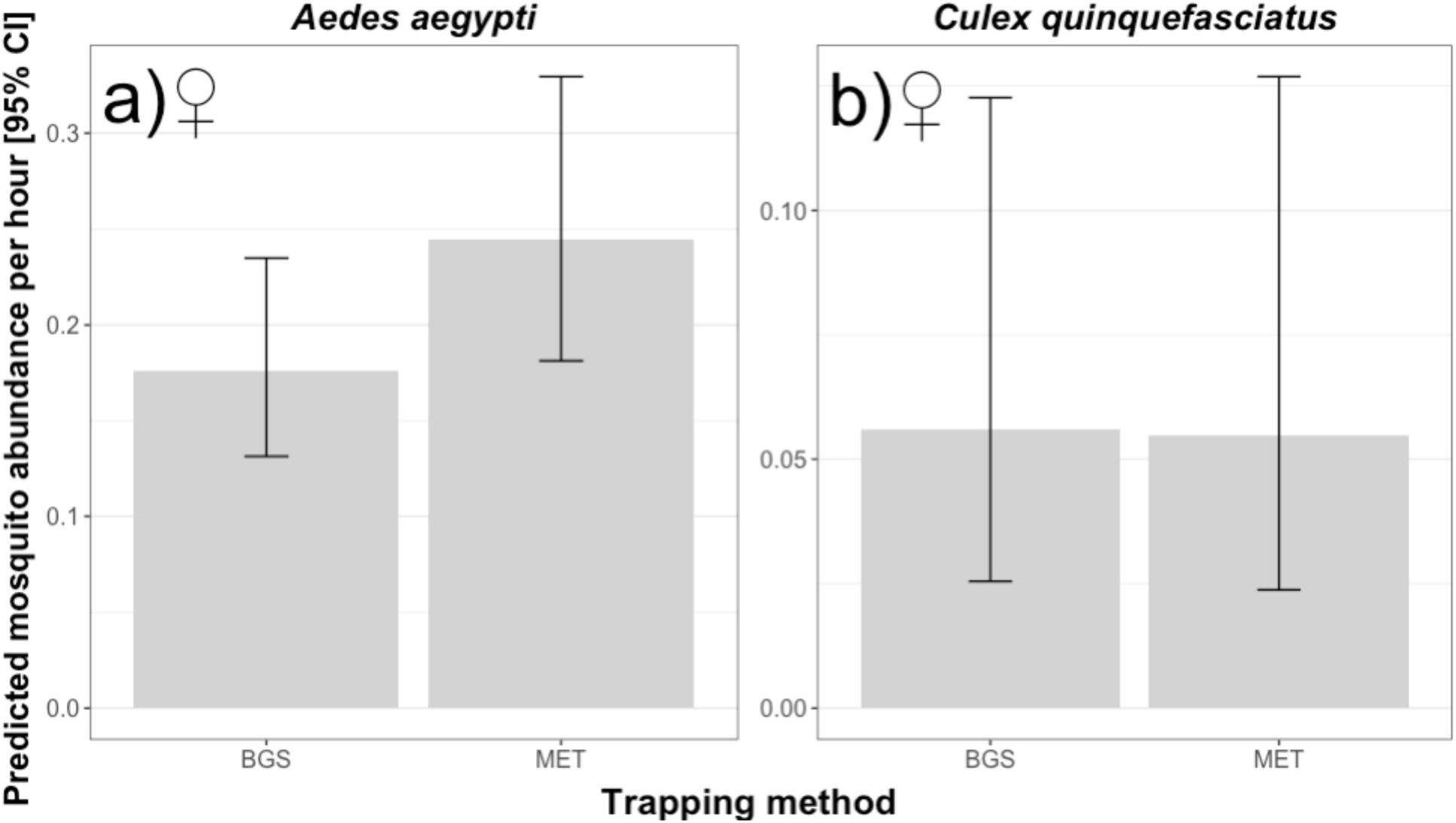
Predicted hourly abundance of mosquitoes using different trapping methods. Panel a) represents *Ae. aegypti* and b) *Cx. quinquefasciatus*. The error bars indicate the Confidence Intervals (C.I.) at 95%.

**Figure 7.**
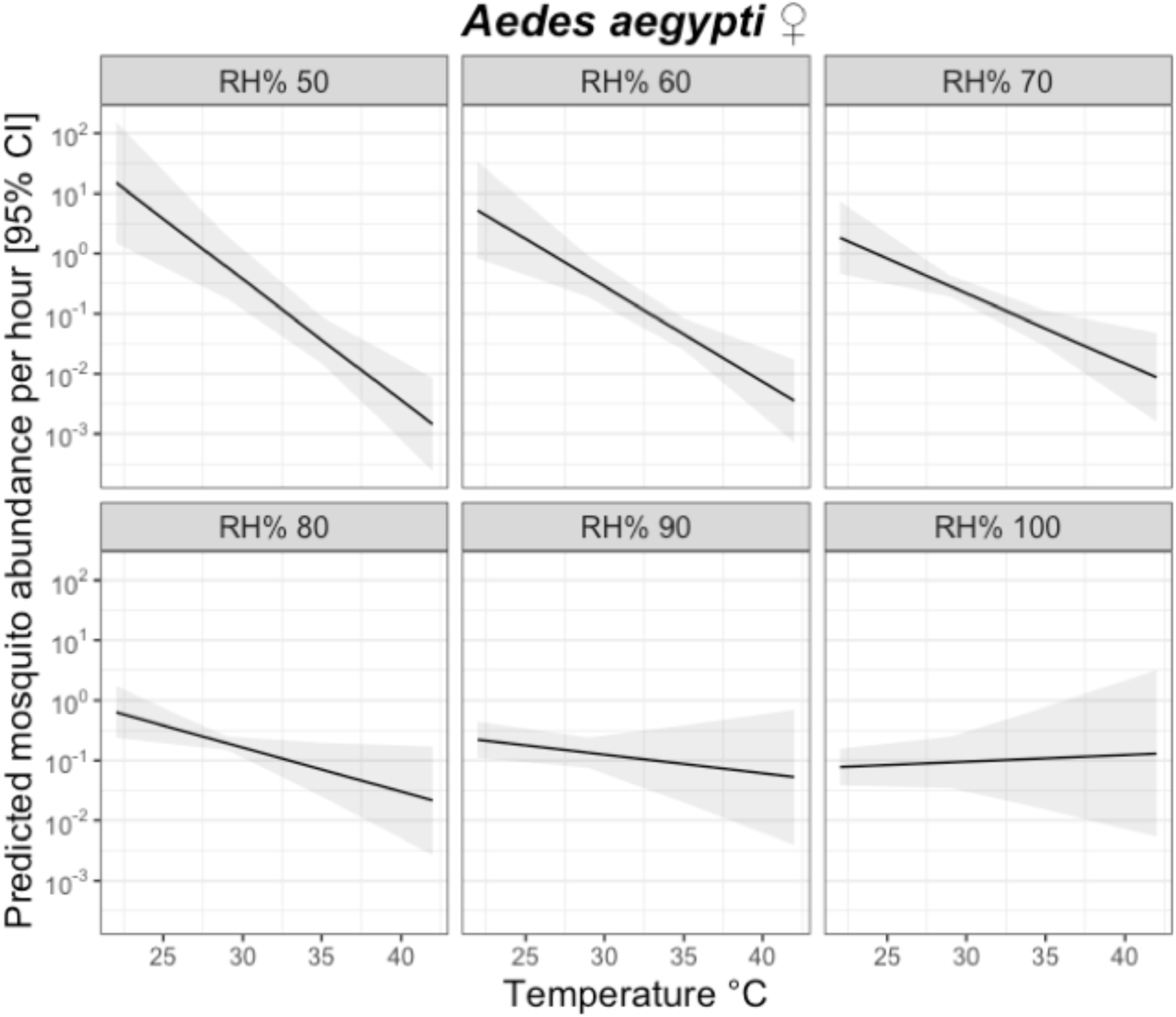
Predicted relationship between the hourly abundance of *Ae. aegypti* females and mean temperature (°C) under different relative humidity (RH) conditions. The black line represents the predicted abundance of *Ae. aegypti* in that hour, with the shaded area representing the 95% Confidence Intervals (C.I.).

The biting activity of female *Cx. quinquefasciatus* also varied significantly across the sampling day. As with *Ae. aegypti*, this pattern was characterized as a quadratic relationship in which mosquito activity peaked during the early morning and late afternoon (Table 3, Figure 5b). Accounting for this activity pattern, there was no difference in the number of *Cx. quinquefasciatus* caught per hour in different trapping methods (Table 3, Figure 6b), and no association with temperature or humidity.

### Molecular screen for ZIKV, DENV and CHIKV

All positive controls worked in all of the PCRs carried out in this study. PCRs on S7 showed that RNA was successfully transcribed into cDNA in 121 mosquito pools out of 122 (Figure S2). Weak bands coming from a few samples were observed on the gels from ZIKV, DENV 1-3 (PCR from universal primer) and DENV 4 (Figure S3, Figure S4 and Figure S5). Then, a second PCR was run on only those samples for reconfirmation. Weak bands were observed for a second time in some of the samples of ZIKV (Figure S6a). However, a PCR on the negative RT of those samples also showed weak bands (S6b) suggesting that they may be due to contamination with gDNA. Positivity of DENV 4 could not be confirmed either as no bands were observed on any of the samples (Figure S7). The second PCR of DENV 1-3 showed weak bands for the second time on a few samples (Figure S8), which were then tested on three separated PCRs using individual primers of DENV1, DENV2 and DENV3. However, positivity could not be confirmed either on any of the samples as no bands were observed (Figure S9). PCR results confirmed that no samples were infected with CHIKV either (Figure S10).

## DISCUSSION

Identifying an accurate method to predict the exposure of humans to infected mosquito vectors has been an enormous challenge for *Aedes*-borne pathogens [81,100]. Here, we present the MET as a potential alternative for safe measurement of *Aedes* landing rates on humans. When tested in dengue, chikungunya and Zika-endemic settings in Ecuador, the MET provided similar estimates of *Ae. aegypti* abundance and biting activity as the current gold standard BGS sentinel method. While the BGS uses artificial odour baits and carbon dioxide (CO_2_) to lure mosquitoes into a standardized trap; the MET directly estimates the number of *Aedes* host-seeking within the immediate vicinity of a real host. The standardization provided by the BGS makes it easy and effective to use in widescale surveillance [82,84], although a limitation is that non-biogenic CO_2_ sources are not always available [71,78]. However, the degree to which BGS collections accurately reflect *per capita* human biting rates is unclear. For example, BGS trapping efficiency may vary with the type and number and types of lures used, and volume of C0_2_ released [60,76,101], making it difficult to infer how they translate to exposure experienced by one person in that environment. An advantage of the MET is that it is more directly analogous to the human landing catch in sampling mosquitoes in the process of host seeking on a person. This could also be seen on the total catches of the other mosquito species when compared to the total numbers trapped by the BGS. The MET could thus provide a useful supplementary surveillance method for estimation and validation of human biting rates and the associated entomological inoculation rate (EIR).

By facilitating a safe and more direct estimation of the EIR for *Aedes*-borne viruses, the MET could provide robust and precise entomological indicators of transmission intensity [85–87]. Such indicators are much needed to understand heterogeneity in transmission [3,4,55], and evaluate the efficiency of vector control interventions. However this relies on the assumption that the MET accurately reflects the true *Aedes* exposure of one person per unit of time. Estimates of human exposure to the malaria vector *An. gambiae s.l.* from the MET were similar to those of the human landing catch in some studies [87,102]; whereas in others mosquito abundance was underestimated by the MET compared to the HLC [86]. Here it was not possible to directly compare the MET to the HLC because of ethical restrictions in using the latter in an area of high arboviral transmission. However we speculate that one factor that could cause the MET to underestimate *Aedes* vectors biting rates is the area of the body protected. Whereas African *Anopheles* vectors generally prefer feeding on the lower legs and feet [103–105]; it is not clear if *Aedes* prefer to bite on specific parts of the body [106,107]. As a next step in validating this approach, we recommend the MET to be directly compared to the HLC under controlled conditions with uninfected *Aedes* vectors (*e.g.* semi-field experiments).

Both the MET and BGS trap sampled a similar composition of mosquito species in the study period. However, estimates of the mean daily and hourly abundance of *Ae. aegypti* and *Cx. quinquefasciatus* were slightly but not statistically higher in MET than BGS collections. The relatively short period of this (12 sampling days) may have limited power to detect for minor-moderate differences between trapping methods. We thus conclude the MET is at least as good as the BGS gold standard for sampling host-seeking *Aedes* vectors in this setting, but also recommend further longer-term comparisons over a wider range of seasons, sites and participants to evaluate whether the MET outperforms the BGS. If we assume that MET is equivalent to HLC, these results are also consistent to those shown by Kröckel *et al.*, who also observed that HLC captured more mosquitoes, although not statistically different from the BGS [84].

Mosquito collections conducted here were also used to test for associations between *Aedes* host-seeking activity and microclimatic conditions. The impact of temperature and humidity on the life-history, physiology, behaviour and ecology of *Ae. aegypti* has been extensively investigated under laboratory conditions [108–116]. However, relatively little is known about how microclimate impacts the diel host-seeking behaviour of wild *Aedes*. In general, the host-seeking activity *Ae. aegypti* and *Cx. quinquefasciatus* was higher on days when mean temperatures were lower (across range of 25°C to 30°C). Additionally, the hourly biting rates of *Aedes* were negatively associated with temperature but only under conditions of low humidity. As mean hourly temperatures were strongly negatively correlated with relative humidity (Figure S1), these results indicate that *Ae. aegypti* biting activity is highest during relatively cool and humid hours of the day. These microclimatic associations may account for the observed biting activity of *Ae. aegypti* and *Cx. quinquefasciatus*. A comprehensive review [99] of *Ae. aegypti* biting behaviour indicates that bimodal and trimodal activity patterns are often reported, with evidence of specific adaptations to other ecological features (*e.g.* artificial light availability) [99]. Such variability seems to be common and related to optimal humidity and temperature conditions available during such hours [117,118].

A key feature of any method for estimating EIR is its ability to estimate human biting rates and infection rates in mosquitoes. While the results here presented indicate that the MET could be used to estimate the human biting rates, the infection rates could not be measured as none of the *Aedes* mosquitoes collected with either trapping method were positive for arboviruses. Reported rates of arboviruses in *Aedes* vectors are generally very low (0.1% to 10%) even in high transmission areas (*e.g.* [119–126]). Thus failure to detect arboviruses within the relatively small sample size of vectors tested here (*e.g.* 207 individuals tested in 122 pools) is not unexpected.

Although promising, the MET has a number of limitations relative to the BGS for sampling host seeking *Aedes*. First, it requires human participants and the trap itself is heavier, which is more labour intensive than using BGS. Also, as the METs used here are still research prototypes produced on a bespoke basis without a licenced manufacturer, their production cost is currently more expensive than BGS traps (approximately £650 versus £170 per trap, respectively). In addition, some technical problems were experienced including a tendency to short circuit under conditions of high air humidity. These limitations are expected to be improved if manufactured at scale as manufacturing costs would fall and technical improvements should make the MET suitable for humid environments. The enormous advantage of the MET is therefore, its potential ability to estimate the EIR for arboviral infections.

## CONCLUSIONS

Here we evaluated the MET as a tool for estimating human biting rates of the arboviral vector *Ae. aegypti* in a high transmission setting in coastal Ecuador. The MET performed at least as good as the current BG-sentinel trap gold standard for estimating the mean abundance per hour of host-seeking *Aedes*, and provided a realistic representation of hourly activity patterns. We conclude MET is a promising tool for *Ae. aegypti* and other mosquito species surveillance, which could uniquely enable a relatively direct estimate of the arboviral entomological inoculation rate experienced by communities.

## DECLARATIONS

### Ethical approval and consent to participate

Ethical approval for this research was granted by the MVLS College Ethics Committee of the University of Glasgow (Project No.: 200150175), and by the Ethics Committee of Research on Human Beings of the San Francisco de Quito University (2016-146M).

Prior to the study, the objectives of this research and risks and benefits for taking part in this were explained to participants in MET collections and their written informed consent was obtained. Oral informed consent was also obtained from the head of households where mosquito collections were performed. The purpose and objectives of the study was explained to householders before requesting permission for the study team to collect mosquitoes within their property.

### Agreements and permits for conducting research

The Government of Ecuador, through the Ministry of Environment (MAE), granted the permits to carry out the present study under the Framework Agreement on Access to Genetic Resources No. MAE-DNB-CM-2016-0052. Transportation of samples from the study site in Quinindé (Rosa Zárate) to the LEMMT-USFQ in Quito was authorized by MAE through the document No. MAE-DPAE-2017-1163-O. Transfer of samples from the LEMMT-USFQ in Ecuador to the MRC-University of Glasgow CVR in the United Kingdom, was firstly established by a Material Transfer Agreement signed between Universidad San Francisco de Quito and the University of Glasgow on the 1^st^ of September of 2017. The exportation of biological samples from Ecuador was authorized by MAE under the document 076-17-EXP-IC-FAU-DNB/MA. Finally, a certification of no-need of importation permit to Scotland was issued through a letter from the Animal Health and Welfare Division of the Agriculture and Rural Economy Directorate of the Scottish Government on the 14^th^ of September of 2017.

### Consent for publication

Individuals pictured in the photographs of this manuscript granted their oral consent of the photographs in which they appear to be published.

### Availability of data and materials

The dataset generated and analysed during this study is publicly available in the Open Science Framework repository at https://osf.io/zwbs8.

### Competing interests

Authors declare there are no competing interests.

### Funding

Funding for LDOL to develop this research as part of his PhD studies was provided by the Government of Ecuador through SENESCYT (AR2Q-9554) under the programme “Primera Convocatoria Abierta 2013”. Funding for this research work was provided by the Medical Research Council from the United Kingdom under the grant number MC_PC_15081 (AK, HMF, LDOL, RL) and MC_UU_12014/8 (EP, AK).

### Authors contributions

LDOL, HMF, RL and AK conceptualized the project. LDOL, MPB, ST and SS conducted the field work of mosquito collections under the supervision of RL and HMF. MPB and LDOL carried out the morphological analyses. LDOL and FA carried out the molecular analyses under permanent supervision of EP and AK. LDOL carried out the statistical analyses and wrote the manuscript under the guidance of HMF. AK, EP, HMF, MPB and RL edited the manuscript. All authors read and approved the final manuscript.

## Acknowledgements

We thank all the residents from the local communities for granting access to their properties every day of the study. We also want to thank Jakub Czyzewski and the rest of the Bioelectronics Unit team of the University of Glasgow for building the Mosquito Electrocuting Traps (METs) and providing with their technical assistance before and during the experiments. We also want to specially thank Miguel Ortega-López and Teresa López-Cuesta for designing and building the retractable aluminium frames and the mosquito netting for the METs. We finally thank Ana Espinoza from Fibios Science Communication (www.fibios.org) for the design of the graphical abstract of this manuscript.

## Notes

https://osf.io/zwbs8

