## Supplementary methods and sup. figures for "The Mosquito Electrocuting Trap As An Exposure-Free Method For Measuring Human Biting Rates By *Aedes* Mosquito Vectors"

### Supplementary Materials and Methods

#### *RNA extraction, reverse-transcription and PCR*

Mosquito pools were placed in 1.5 ml crio-vials containing TRIzol™ (Invitrogen) and stored at -80°C for RNA preservation at LEMMT-USFQ before being shipped on dry ice to the MRC-University of Glasgow Centre for Virus Research (CVR) for further analysis. At CVR, samples were transferred to a -80 °C freezer until further processing.

To extract RNA, glass beads were added to each mosquito pool and used to homogenize the sample with a Precellys 24™ homogenizer (Bertin Instruments; homogenization for 30 seconds at 6800 rcf in TRIzol™ Reagent). RNA was then extracted from samples using TRIzol™ according to the manufacturer's instructions; with the exception that 1-Bromo-3-chloropropane (Sigma) being used instead of chloroform. RNA was precipitated with isopropanol and the pellet washed with 75% ethanol and then re-suspended in 50 µl of nuclease-free water. The concentration and quality of the extracted RNA was measured using a NanoDrop 2000™ (Thermo Fisher Scientific). RNA samples were subsequently aliquoted and stored at -80 °C until the reverse-transcription (RT) step.

RTs as well as negative RTs were performed using Moloney Murine Leukemia Virus Reverse Transcriptase™ (M-MLV RT) (Promega) from 950.4 ng of RNA in 40 µl final according to the manufacturer's instructions. cDNAs were aliquoted and further stored at -20°C until PCR was carried out. Out of 147 pools, the concentration of RNA in 25 was too low for analysis and thus excluded from further processing the analysis.

PCR was performed on cDNA prepared from mosquito pools using the DreamTaq Green Polymerase (ThermoFisher) following manufacturer's instructions. A first round of PCR was run on all samples to amplify the S7 gene (ribosomal protein) to assess the RNA integrity of the sample (*i.e.* RNA successfully reverse-transcribed into cDNA). For all samples where RT was confirmed to have succeeded, subsequent PCRs were performed to amplify specific cDNA regions to detect ZIKV, DENV (1-4 strains) and CHIKV. Primers, melting temperatures and expected sizes used are listed in Table S1. Some published primers were shortened to have the same melting temperature for both forward and reverse primers. To detect the reference gene S7 (ribosomal protein, 35 PCR cycles) or arboviruses (40 PCR cycles), PCR samples were respectively run on 1.5% or 3% agarose gel containing EtBr and visualized with a GelDoc imaging system (Biorad). DNA ladders used for 1.5% and 3% gels were respectively Gene Ruler 100 bp Plus (Thermo Scientific) and Ultra Low Range DNA ladder (Invitrogen).

Primers used for this study were previously designed and used for virus screening [1–4]. In addition, we checked that they matched relevant strains for each arbovirus to minimize the possibility of missing an infected sample. PCRs for ZIKV, DENV and CHIKV were performed on all of the samples that were sucessfully transcribed into cDNA.

cDNAs from experimentally ZIKV-infected *Ae. aegypti* females (infectious blood meal at $10^7$ PFU/mL, whole females 14 days post infection, ZIKV clone pCCI-SP6-ZIKV-Rz [5]) were used as a positive control for ZIKV. For DENV1 and DENV1-3, cDNA from a confirmed infected mosquito with DENV1 was used as a positive control (different field collection, unpublished data). For the rest of the viruses, regions corresponding to the amplified sequence were designed based on previously published virus isolates from GenBank and synthetized (GENEWIZ), diluted at 3 ng/µL and used as positive controls (Table S2).

**Table S1. Primers used for detection of arboviruses by RT-PCR.** Assays included S7 (ribosomal protein) for RNA integrity, Zika virus (ZIKV), universal assay for dengue virus strains 1, 2 and 3 (DENV 1-3), dengue virus strain 1 (DENV1), dengue virus strain 2 (DENV2), dengue virus strain 3 (DENV3), dengue virus strain 4 (DENV4), and chikungunya virus (CHIKV).

| Assay | Primer name | Sequence (5'→3') | Reference | Expected size (bp) | Melting temperature |
|---|---|---|---|---|---|
| S7 | S7 (forward) | GGGACAAATCGGCCAGGCTATC | Newly designed | 403 with intron 290 without | 58°C |
| | S7 (reverse) | TCGTGGACGCTTCTGCTTGTTG | | | |
| ZIKV | Zika 1087 (forward) | CCGCTGCCCAACACAAG | Modified from [1] | 76 | 57°C |
| | Zika 1163c short (reverse) | CCACTAACGTTCTTTTGCAG | | | |
| DENV 1-3 | DENV_F (forward) | GCATATTGACGCTGGGARAGAC | [2] | 63 | 67°C |

| | | | | | |
|---|---|---|---|---|---|
| | DENV_R1-3 | TTCTGTGCCTGGAATGATGCTG | | | |
| DENV1 | DENV1_F (forward) | CAATGGATGACAACAGAAGAYATG | [3] | 71 | 60°C |
| | DENV1_R (reverse) | TCCATCCATGGGTTTTCCTCTAT | | | |
| DENV2 | DENV2_F (forward) | GCAGAAACACAACATGGAACRATAGT | | 199 | 60°C |
| | DENV_2R (reverse) | TGATGTAGCTGTCTCCRAATGG | | | |
| DENV3 | DENV3_F (forward) | ATGGAATGTGTGGGAGGTGG | | 167 | 60°C |
| | DENV3_R (reverse) | GGCTTTCTATCCARTAGCCCATG | | | |
| DENV4 | DENV_F (forward) | GCATATTGACGCTGGGARAGAC | [2] | 63 | 60°C |

| | | | | | |
|---|---|---|---|---|---|
| | DENV_R4 (reverse) | YTCTGTGCCTGGATWGATGTTG | | | |
| CHIKV | CHIKF short (forward) | ACCGGCGTCTACCCATT | Modified from [4] | 312 | 56°C |
| | CHIKR short (reverse) | GGGCGGGTAGTCCATGTT | | | |

**Table S2. Positive control DNA sequences used as PCR positive controls.** Sequences synthetized with primer sequences in bold. Sequences were designed according to previously GenBank published virus isolates.

| **Assay** | **Positive control Sequence (5'→3')** | **GenBank sequence used** |
|---|---|---|
| DENV1 | GCCCACCAC**CAATGGATGACAACAGAAGACATG**TTATCAGTGTGGAATAG GGTCTGG**ATAGAGGAAAACCCATGGATGGA**GGACAAAACT | GenBank: KY474307.1 |
| DENV2 | AAGGAAATA**GCAGAAACACAACATGGAACAATAGT**TATCAGAGTACAATA TGAAGGGGACGGTTCTCCATGTAAGATCCCTTTTGAGATAATGGATTTGGA AAAAAGACATGTTTTAGGTCGCCTGATTACAGTCAACCCAATCGTAACAGA AAAAGATAGCCCAGTCAACATAGAAGCAGAACCT**CCATTCGGAGACAGCT ACATCA**TCATAGGAG | GenBank: AF038403.1 |
| DENV3 | CTCAAGAGC**ATGGAATGTGTGGGAGGTGG**AAGATTACGGGTTCGGAGTTT TCACAACCAACATATGGCTGAAACTCCGAGAGGTGTACACCCAACTATGT GACCATAGGCTAATGTCGGCAGCCGTCAAGGATGAGAGGGCCGTACACGC | GenBank: KU050695.1 |

| | | |
|---|---|---|
| | CGA**CATGGGCTATTGGATAGAAAGCC**AAAAGAATG | |
| DENV4 | ACAAAAACA**GCATATTGACGCTGGGAAAGAC**CAGAGATCCTGCTGTCTCTG<br>**CAACATCAATCCAGGCACAGAG**CGCCGCGA | GenBank:<br>AY947539.1 |
| CHIKV | AAGGTCTTC**ACCGGCGTCTACCCATT**CATGTGGGGCGGCGCCTACTGCTTC<br>TGCGACACCGAAAATACGCAATTGAGCGAAGCACATGTGGAGAAGTCCGA<br>ATCATGCAAAACAGAATTTGCATCAGCATACAGGGCTCATACCGCATCCGC<br>ATCAGCTAAGCTCCGCGTCCTTTACCAAGGAAATAATATCACTGTGGCTGC<br>TTATGCAAACGGCGACCATGCCGTCACAGTTAAGGACGCTAAATTCATAGT<br>GGGGCCAATGTCTTCAGCCTGGACACCTTTCGACAATAAAATCGTGGTGTA<br>CAAAGGCGATGTCTAC**AACATGGACTACCCGCCC**TTCGGCGCA | GenBank:<br>KR559470.1 |

**Supplementary Figures**

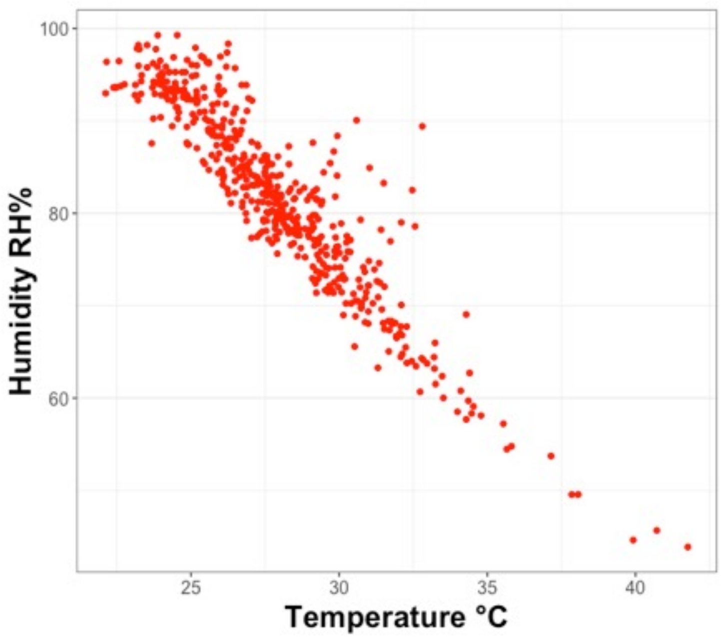


**Figure S1. Relationship between observed Temperature (C°) and Relative Humidity (%).** Red dots represent individual observations per hour recorded.

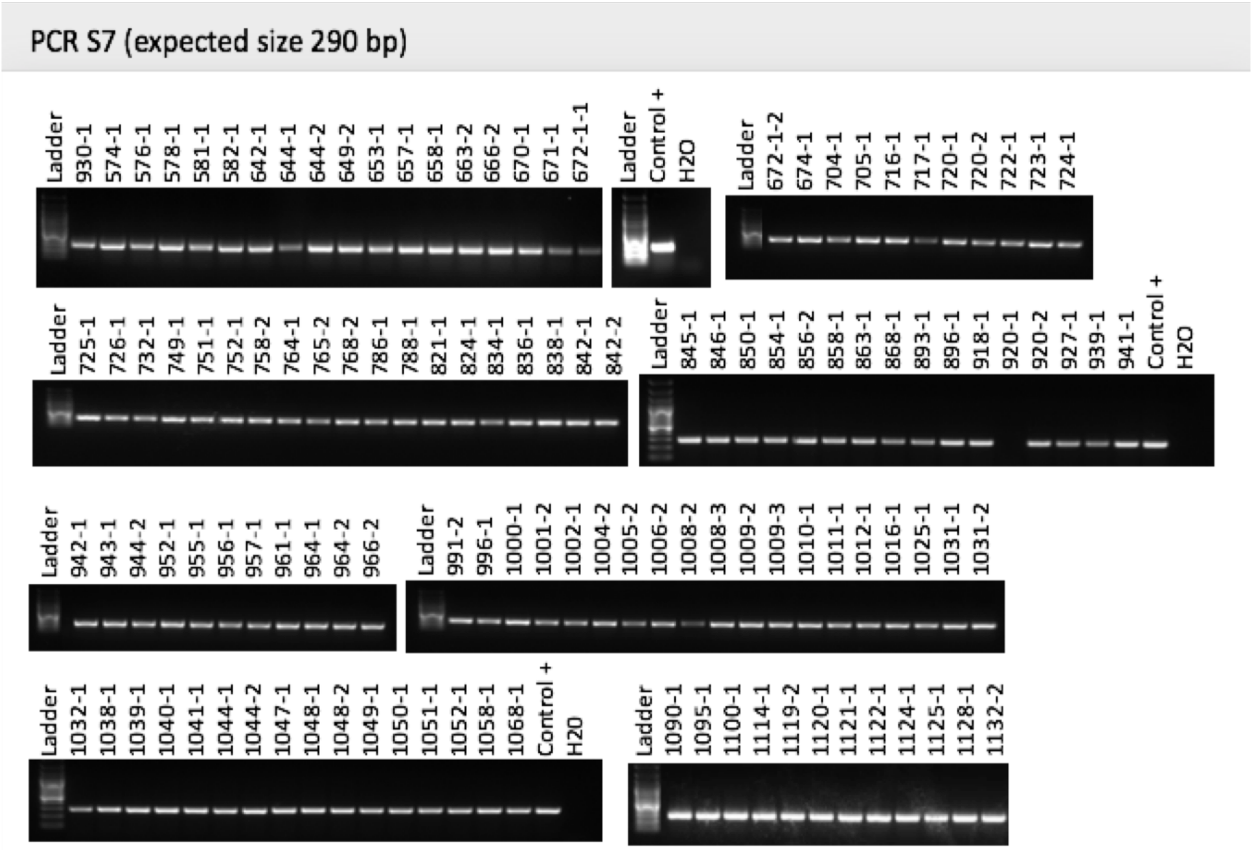


**Figure S2. Visualization of the PCR products of S7 gene on agarose gels.** All of the samples, with the exception of one (sample 920-1), were positive.

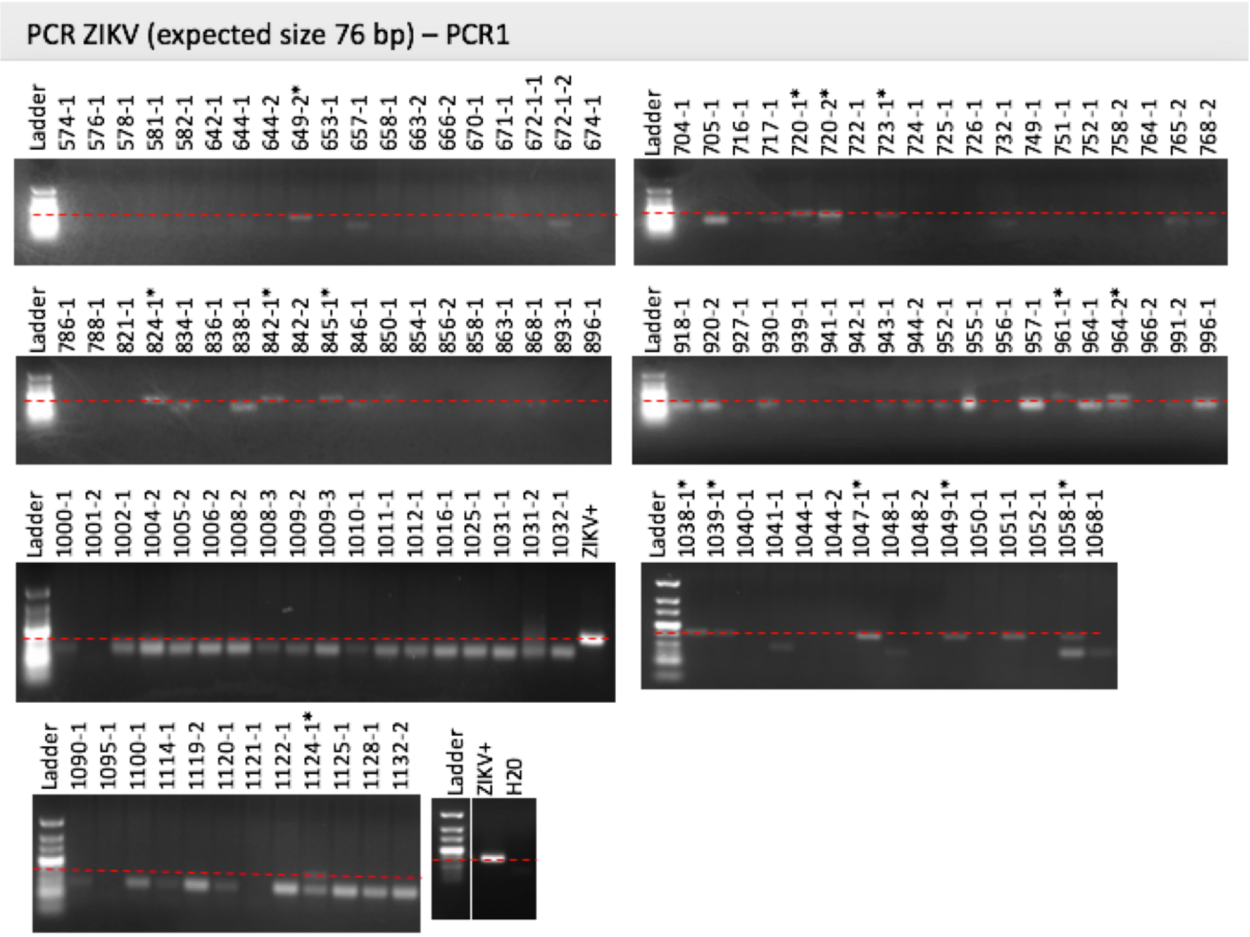


**Figure S3. Visualization of the first PCR products of ZIKV on agarose gels.** The * indicates the samples that had a weak band at the expected size of 76 bp. The red dashed line is positioned at approximately 76 bp. **ZIKV+** corresponds to the positive control.

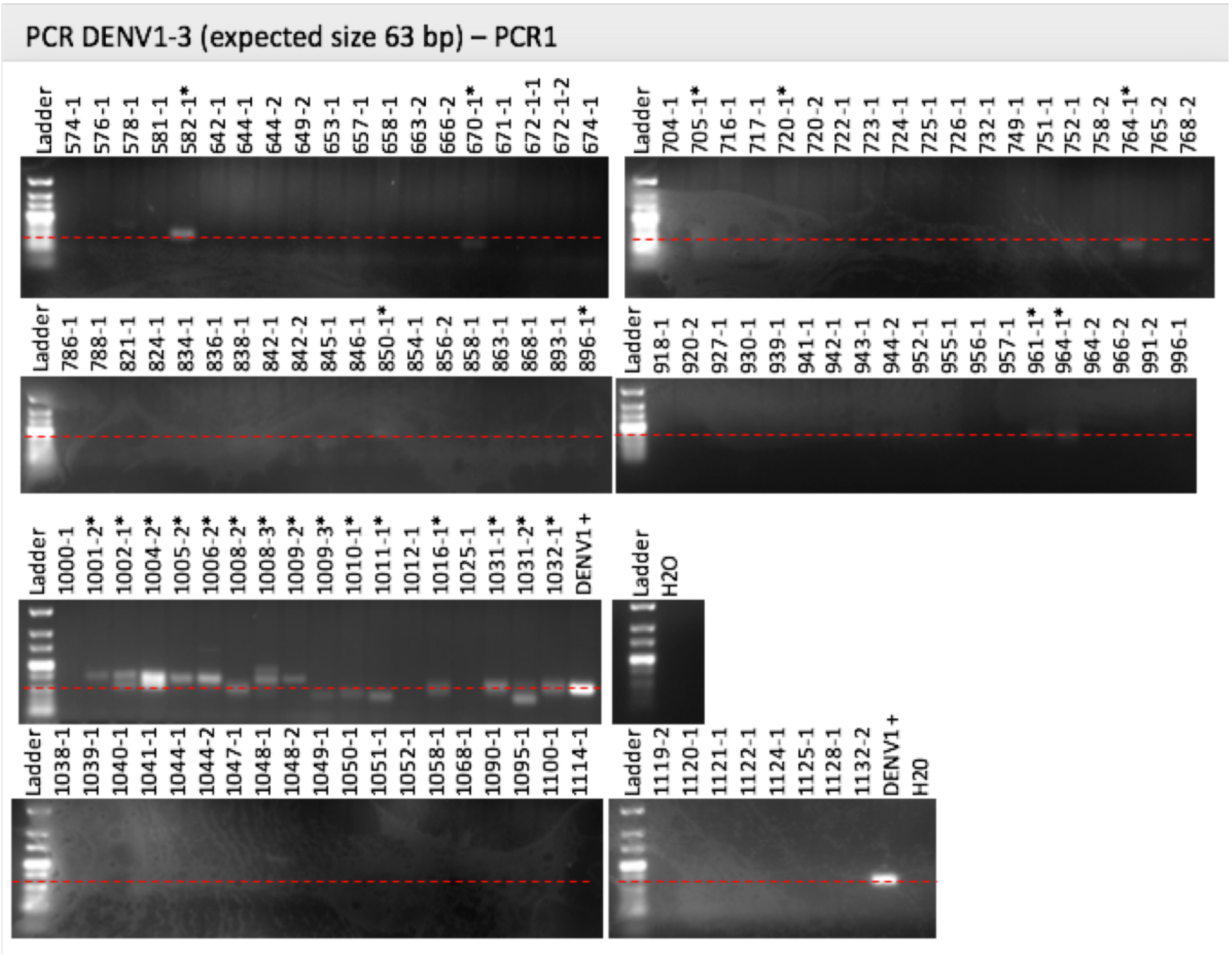


**Figure S4. Visualization of the first PCR products of DENV 1-3 on agarose gels.** The * indicates the samples that had a weak band at the expected size of 63 bp. The red dashed line is positioned at approximately 63 bp. **DENV1+** corresponds to the positive control.

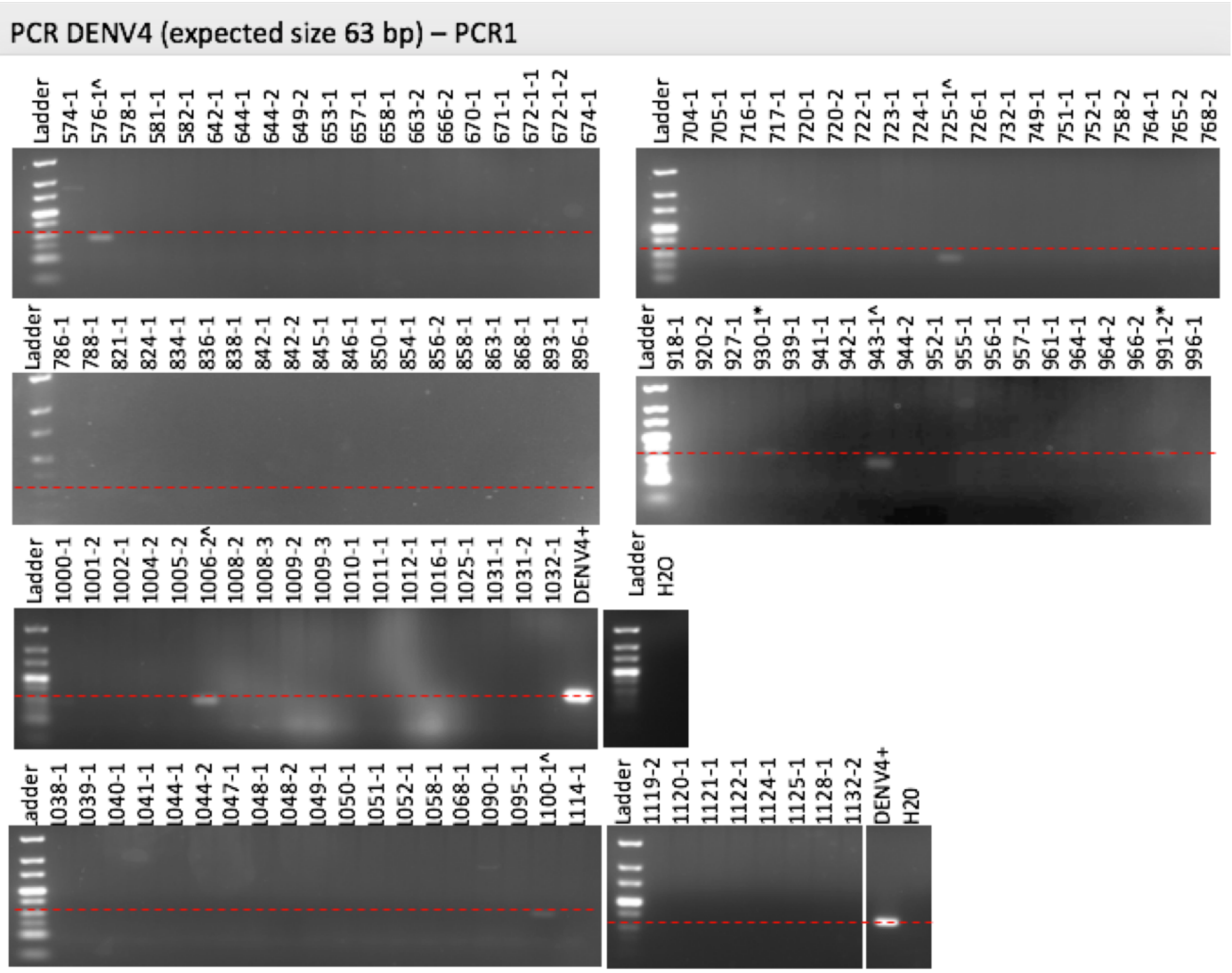


**Figure S5. Visualization of the first PCR products of DENV 4 on agarose gels.** The * indicates the samples that had a weak band at the expected size of 63 bp and **^** indicates the samples that showed a size close to the expected one. The red dashed line is positioned at approximately 63 bp. **DENV4+** corresponds to the positive control.

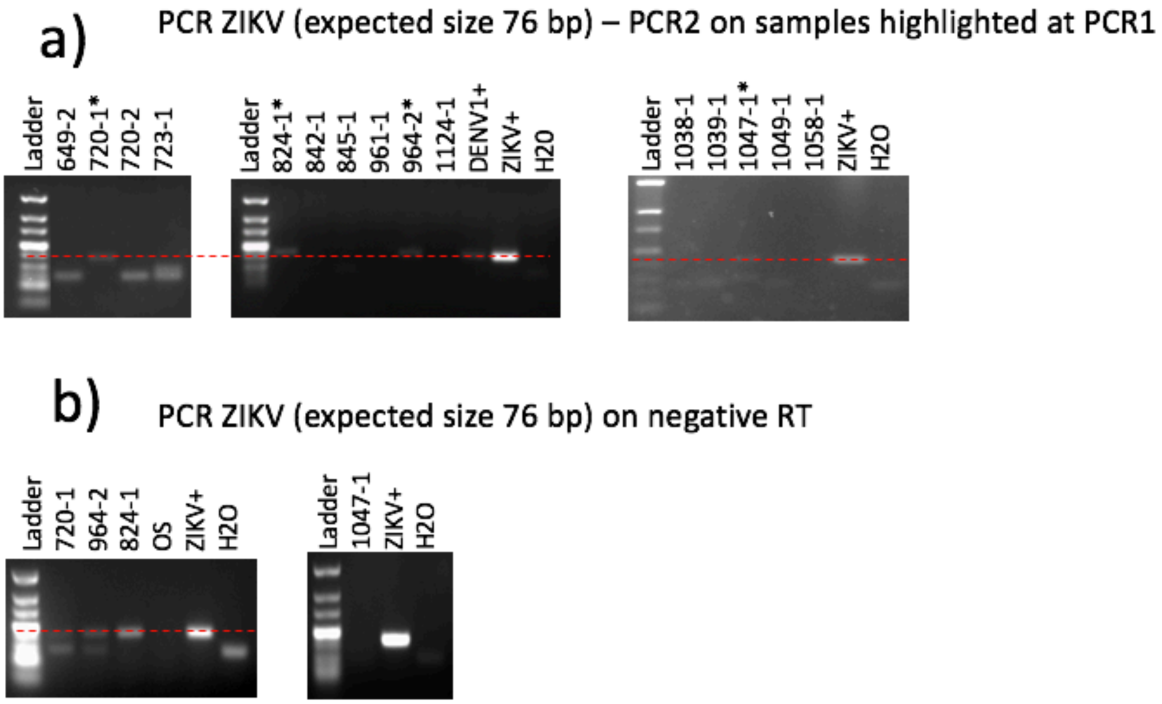


**Figure S6. Visualization of the second PCR products of ZIKV on agarose gels.** The * indicates the samples that had a weak band at the expected size of 76 bp on the first PCR. The red dashed line is positioned at approximately 76 bp. **ZIKV+** is the positive control.

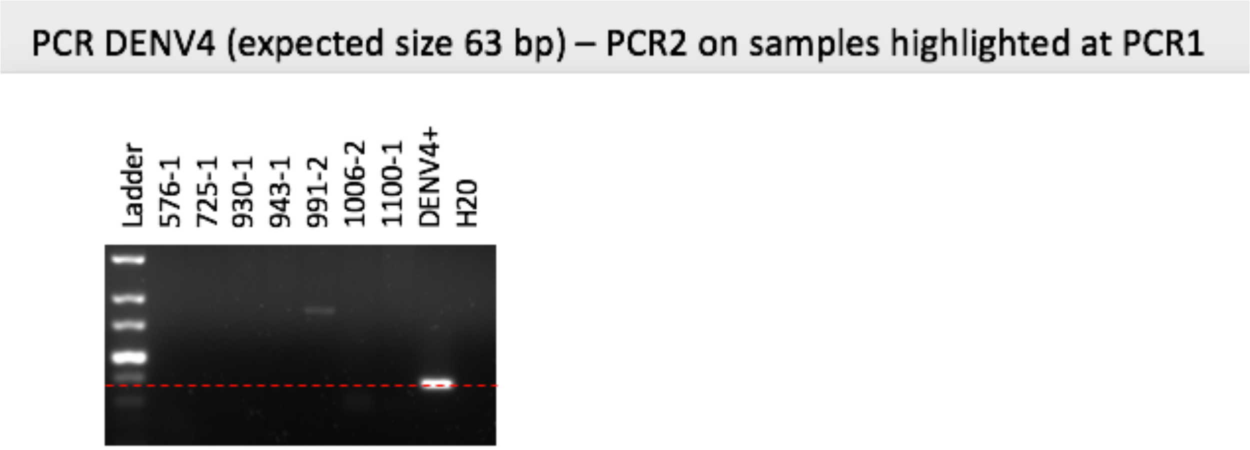


**Figure S7. Visualization of the second PCR products of DENV 4 on agarose gels.** All samples were negative. The red dashed line is approximately positioned at the expected size of 63 bp. **DENV4+** corresponds to the positive control.

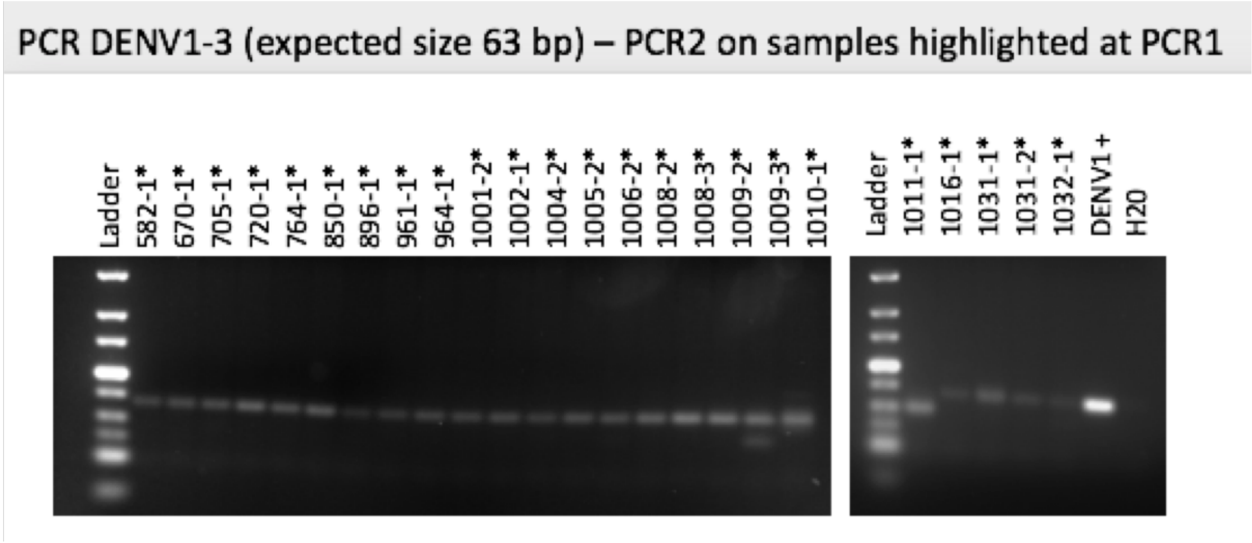


**Figure S8. Visualization of the second PCR products of DENV 1-3 on agarose gels.** The * indicates the samples that a weak was again present on the second PCR at the expected size of 63 bp. **DENV1+** corresponds to the positive control.

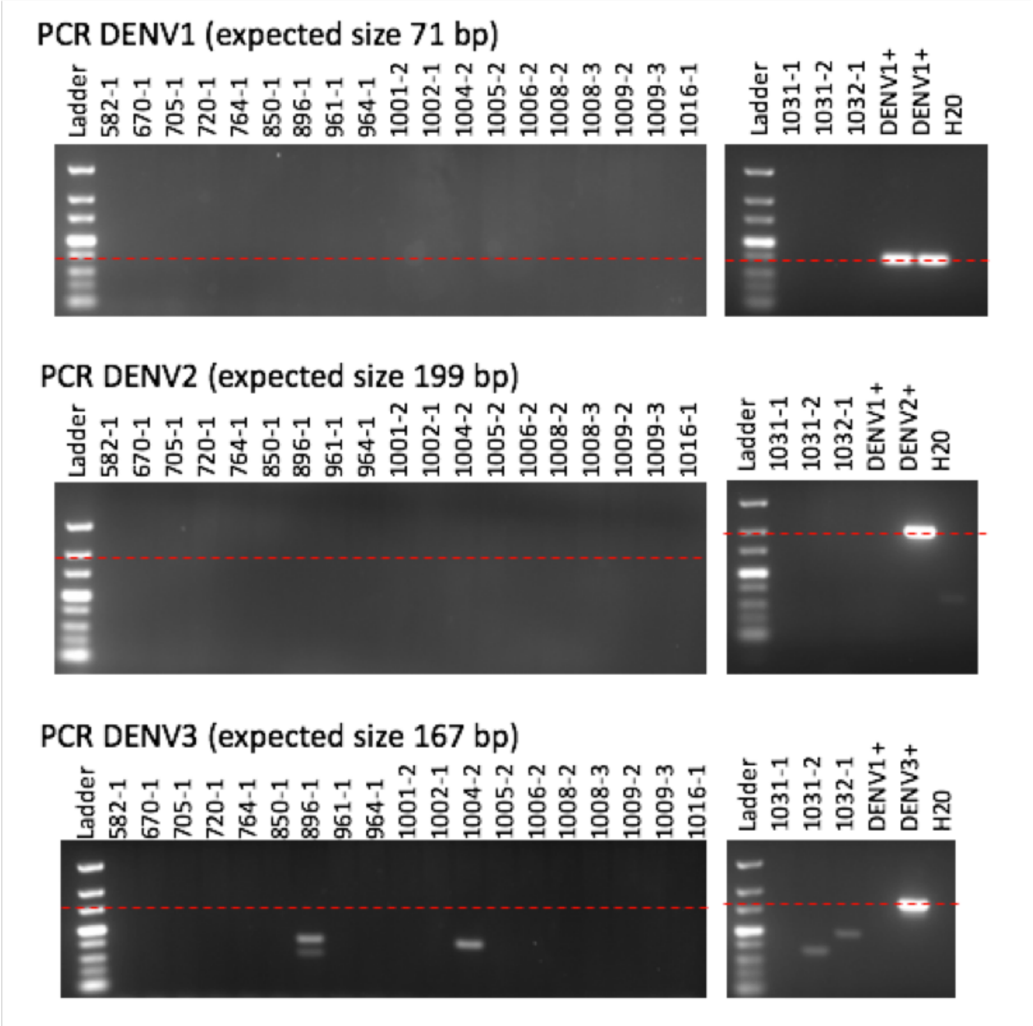


**Figure S9. Visualization of the individual PCR products of DENV1, DENV2 and DENV3.** The same samples from Figure S8 are shown for individual runs for each of the three DENV isotypes. The red dashed line is positioned at approximately 71 bp for the DENV1 run, at 199 bp for the DENV2 run, and at 167 bp for the DENV3 run. **DENV1+, DENV2+** and **DENV3+** correspond to the PCR positive controls.

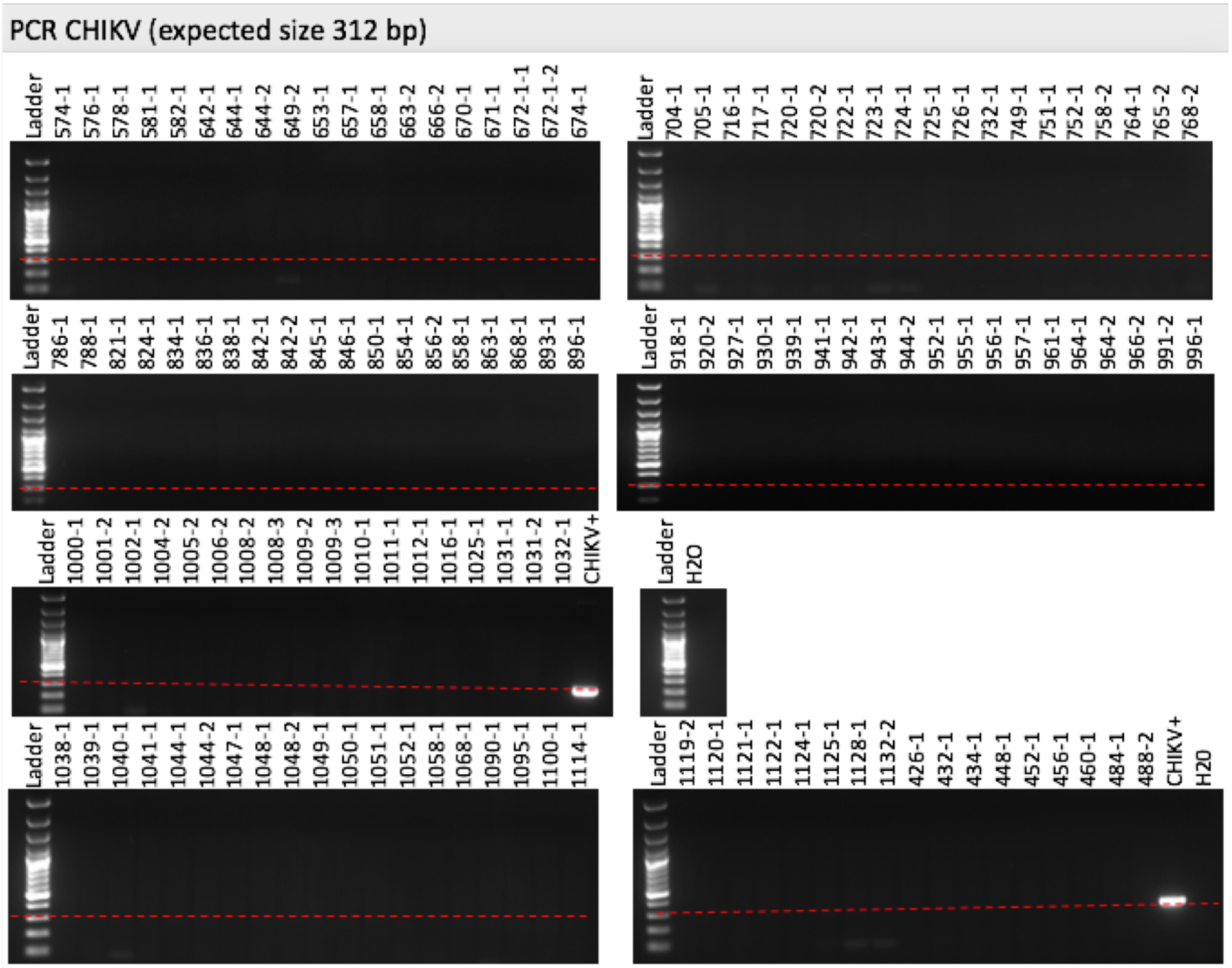


**Figure S10. Visualization of the PCR products of CHIKV on agarose gels.** The red dashed line is positioned at approximately 312 bp. **CHIKV+** corresponds to the positive control.
